# *O*-mannosylation of fibrocystin and fibrocystin-L expands TMEM260 sequon selectivity and provides a potential link to the renal and auditory phenotypes of SHDRA

**DOI:** 10.64898/2026.09.21.753134

**Authors:** Jonas Kaynert, Christina Hopf, Smilla Brand-Merseburger, Christoph Garbers, Andreas Pich, Falk F. R. Buettner

**Author notes:** Corresponding author: Falk F. R. Buettner.

## Abstract

Protein *O-*mannosylation in the endoplasmic reticulum (ER) regulates the maturation and function of extracellular proteins, yet the substrate spectrum and biological functions of individual *O-*mannosyltransferases remain incompletely understood. TMEM260 is an ER-resident *O-*mannosyltransferase that modifies immunoglobulin-like, plexins, transcription factors (IPT) domains. Here, we identify fibrocystin and fibrocystin-L, encoded by *PKHD1* and *PKHD1L1*, respectively, as novel TMEM260-mediated *O-*mannosylated proteins. By combining screening of human proteomic datasets with mass spectrometry-based analysis of IPT domains expressed in TMEM260-deficient HEK293 cells, we demonstrate that multiple IPT domains of fibrocystin and fibrocystin-L are modified by TMEM260. Our findings expand the known substrate requirements of TMEM260 by revealing tolerance of substantial sequon variation and demonstrating serine as an alternative acceptor residue within IPT domains. In humans, loss-of-function variants in TMEM260 cause structural heart defects and renal anomalies (SHDRA) syndrome, whereas pathogenic variants in fibrocystin cause autosomal recessive polycystic kidney disease (ARPKD). Notably, patients with SHDRA and ARPKD share overlapping renal phenotypes, suggesting that impaired TMEM260 function may phenocopy aspects of fibrocystin-associated diseases. Given that fibrocystin is a substrate protein for TMEM260-mediated *O-*mannosylation, these findings raise the possibility that functional impairment of TMEM260 causes aberrant *O-*mannosylation of fibrocystin. Thereby fibrocystin function might be compromised leading to the renal phenotype observed in patients with TMEM260 mutations. To directly assess the functional relevance of this modification, we expressed recombinant fibrocystin fragments in TMEM260-deficient cells complemented with either wild-type TMEM260 or a catalytically inactive TMEM260 mutant. Further, we investigate the *O-*mannosylation of the threonine residue directly *C-*terminal to the ARPKD-associated patient mutation L1407R. Strikingly, substitution of leucine L1407 by the patient-associated arginine residue, completely abolished *O-*mannosylation of the adjacent threonine, demonstrating that this disease-associated mutation directly disrupts a TMEM260-dependent *O-*mannosylation site. Together, our findings establish fibrocystin and fibrocystin-L as TMEM260-dependent *O-*mannosylated proteins and reveal a potential molecular mechanism linking defective TMEM260-mediated *O-*mannosylation to the renal phenotype of SHDRA, which overlaps with *PKHD1*-associated ARPKD.

## Introduction

To date, three *O-*mannosylation pathways have been described in vertebrates, mediated by POMT1/POMT2 (1), members of the TMTC protein family (2) and, most recently, TMEM260 (3). TMEM260 is an endoplasmic reticulum (ER)-resident glycosyltransferase that mediates a distinct *O-*mannosylation pathway targeting IPT (Ig-like, plexins, transcription factors) domains present in plexins and the receptor tyrosine kinases c-MET and RON (3). Despite its restriction to IPT domain bearing proteins, TMEM260 does not modify all IPT domains, indicating substantial substrate selectivity. This selectivity is governed by a sequence motif located within the first two β-strands of IPT domains. Recent structural and sequence analyses suggested that the luminal region of TMEM260 recognizes a degenerated sequon (Px_6_Px_3_Px_2_GGx_3_T), positioning the acceptor threonine residue within the catalytic site. In contrast to canonical glycosylation sequons for *N*- and *C*-mannosylation, this motif tolerates variability at conserved positions, limiting its predictive power and raising the possibility that additional unidentified IPT domain bearing proteins are substrates of TMEM260. (4)

Genetic defects in TMEM260 cause structural heart defects and renal anomalies (SHDRA) syndrome, a multisystem disorder characterized by prominent congenital cardiac and renal abnormalities. Among the cardiac manifestations, persistent truncus arteriosus (PTA) and ventricular septal defects (VSD) are particularly prevalent (5). Notably, highly similar cardiac phenotypes have been reported in patients carrying pathogenic variants of *PLXND1*, which encodes the transmembrane guidance receptor plexin-D1 (3, 5, 6). Plexin-D1 contains multiple IPT domains that are subject to *O-*mannosylation by TMEM260, and the striking phenotypic overlap between TMEM260- and PLXND1-associated disease has therefore suggested that impaired *O-*mannosylation of plexin-D1 may underlie, at least in part, the cardiac abnormalities observed in SHDRA (3, 5, 6). This phenotypic and molecular connection provides a compelling example of how disruption of TMEM260-dependent *O-*mannosylation can translate into congenital developmental defects.

In contrast, in patients with genetic defects in TMEM260 the molecular basis of the renal phenotype remains unresolved. Fibrocystin, a 446 kDa protein encoded by the *PKHD1* gene, and its paralog fibrocystin-L (*PKHD1L1*) contain 12 and 14 IPT domains, respectively, that do not conform to the currently defined TMEM260 sequon requirements (7, 8). Interestingly, pathogenic variants in *PKHD1* cause autosomal recessive polycystic kidney disease (ARPKD), characterized by progressive renal cyst formation (9). However, the *O-*mannosylation of both fibrocystin and fibrocystin-L is unexplored, leaving it open whether altered fibrocystin *O-*mannosylation contributes to the renal phenotype observed in SHDRA.

Here, we first interrogate publicly available human mass spectrometry datasets to identify *O-*mannosylated peptides derived from the IPT domains of fibrocystin and fibrocystin-L. This analysis revealed *O-*mannosylation within the IPT domains of both proteins, suggesting that they may represent previously unrecognized substrates of the TMEM260-dependent *O-*mannosylation pathway. We subsequently validated and extended these findings using TMEM260 knockout (KO) human embryonic kidney 293 (HEK293) cells, complemented by recombinant expression of wild-type and functionally inactive TMEM260. We expressed soluble IPT repeats of fibrocystin and fibrocystin-L in these cells and analyzed *O-*mannosylation upon purification by LC-MS/MS analysis of candidate peptides. Finally, we investigated the ARPKD-associated fibrocystin missense variant L1407R within the IPT9 domain (8), which affects a conserved residue adjacent to a TMEM260 *O-*mannosylation site, to determine whether this disease-associated mutation disrupts *O-*mannosylation and might thereby provide a potential molecular link to ARPKD pathogenesis.

## Materials and methods

### Cell culture

HEK293 cells were grown in high glucose (4.5 g/L) Dulbecco’s Modified Eagle’s Medium (DMEM) supplemented with 10% FCS.

### Cloning procedures

The respective IPT coding sequences were synthesized as gene fragments (TWIST Bioscience, San Francisco, CA, USA) and cloned into the multicloning site of a pcDNA3.1 vector (Thermo Fisher Scientific, Waltham, MA, USA). Each construct included a *N*-terminal secretion peptide (from human C6 complement protein) and a *C*-terminal ALFA (10) - and His10-tag. All plasmids used in this study are listed in table S1.

### CRISPR-Cas9 mediated generation of TMEM260-KO cells

Single guide RNA (sgRNA) sequence (20 nt, see table S2) targeting TMEM260 was designed using CCTop (11). A sgRNA targeting distinct protospacer region in exon 3 was selected from CCTop results for KO generation. Complementary oligonucleotides were synthesized (Sigma-Aldrich, St. Louis, MO, USA), annealed, and ligated into the BbsI site of a gRNA expression vector. For genome editing, 1 µg sgRNA plasmid was co*-*transfected with 1 µg pCas9-GFP into HEK293 cells (6-well format) using PEI MAX® 40,000 (Polysciences, Warrington, PA, USA). GFP-positive cells were sorted 24 h post-transfection by fluorescence-activated cell sorting (FACS) to enrich transfected populations and subsequently expanded as single-cell–derived clones in 96-well plates using culture medium with 20% FBS. Clonal cell lines were screened by PCR amplification and sanger sequencing of the targeted locus. Allelic indel formation in sequencing data was analyzed using TIDE (12). One clone carrying a frameshift mutation introducing a premature stop codon in exon 3 and one wild-type control clone were selected for further analysis and cryopreservation. Clonal genotypes were validated at the transcript level by RNA extraction (TRIzol, Thermo Fisher Scientific), cDNA synthesis (Maxima kit, Thermo Fisher Scientific), and sanger sequencing of the target region (table S2). Primers used for protospacer region amplification are listed in the supplement (table S3).

### In-house production of NbALFA-based tools

The coding sequence of the ALFA-tag specific nanobody (NbALFA) (10) was cloned into a pET24a vector with *C*-terminal His10-tag and transformed into *E. coli* SHuffle® Express competent cells (New England Biolabs, Ipswich, MA, USA) by heat shock. Transformed cells were cultured overnight at 37 °C in LB medium and subsequently expanded in 500 mL Terrific Broth (TB) medium, both supplemented with 50 µg/mL carbenicillin. Protein expression was induced at an OD_600_ of 0.6 - 0.8 by addition of 0.5 mM IPTG, followed by overnight incubation at 30 °C with shaking. Cells were harvested, washed with PBS, and lysed by sonication. The lysate was clarified by centrifugation (27,000 × *g*, 30 min) and sterile-filtered. Soluble NbALFA-His10 was purified from bacterial lysate by immobilized metal affinity chromatography (IMAC) using a 1 mL HisTrap HP column (Cytiva, Marlborough, MA, USA). Further purification and buffer exchange into 0.2 M NaHCO_3_, 0.15 M NaCl (pH 8.3) were performed by size-exclusion chromatography on a Superdex 75 10/300 GL column (Cytiva). Protein purity and molecular weight were verified by SDS-PAGE.

For downstream applications, purified NbALFA was conjugated to IRDye 800CW NHS ester (LI-COR Biosciences, Lincoln, NE, USA), Cy3 NHS ester (manufacturer) and immobilized on NHS-activated Sepharose 4 Fast Flow resin (Cytiva) according to the manufacturers’ instructions. These derivatives were used for western blotting, immunofluorescence and immunoprecipitation, respectively.

### In silico discovery workflow for *O-*mannosylated peptides

Gene symbols for fibrocystin (*PKHD1*) and fibrocystin-L (*PKHD1L1*) were used as search terms in the PeptideAtlas database (13). The database was queried separately for each protein: For fibrocystin, the corresponding PeptideAtlas entry indicated a urinary exosome proteome from healthy individuals with broad coverage of fibrocystin-derived peptides. The associated PRIDE dataset (PXD002279) contained twelve DDA raw files, which were downloaded for subsequent analysis. For fibrocystin-L, the PeptideAtlas entry of a tryptic peptide containing the putative *O-*mannosylation site of IPT2 (PAp02820884) was screened for experiments with the highest number of observations (NObs). This led to the selection of a dataset comprising the surface proteome of different blood cell types (PXD005846). Within this dataset, the peptide of interest was detected only in raw data derived from primary human B cells, T cells, and NK cells. Accordingly, in total six DDA raw files corresponding to these three cell types were downloaded. All raw files were processed using MaxQuant v2.6.4.0 (Max Planck Institute of Biochemistry, Martinsried, Germany) and searched against a custom database containing human IPT domain-containing proteins (listed in Dataset S1). The search was performed using the Andromeda search engine with trypsin/P as the protease, allowing up to 2 missed cleavages. carbamidomethylation of cysteine was specified as a fixed modification, whereas oxidation of methionine, protein *N*-terminal acetylation, and hexose (+162.0528 Da with neutral loss) modification on serine and threonine were set as variable modifications. The precursor mass tolerance was 4.5 ppm after recalibration, and the fragment mass tolerance was 20 ppm. PSMs and proteins were filtered to a 1% false discovery rate (FDR) using a target-decoy strategy. Resulting psm files (msms.txt) were further analyzed in Perseus v2.1.6.0 (Max Planck Institute of Biochemistry) to filter for *O-*mannosylated peptide-spectrum matches (PSMs) derived from fibrocystin and fibrocystin-L. Identified PSMs (see Supplementary Dataset S2) were manually inspected, validated, and MS/MS spectra were visualized using Skyline (v26.1, MacCoss Lab, University of Washington) and FreeStyle (Thermo Fisher Scientific) software packages.

### Recombinant expression and purification of IPT repeats

Cells were grown to 50-70% confluency in 15 cm^2^ dishes and transiently transfected using PEI MAX® 40,000. After 4-6 h, the medium was replaced, and cells and supernatant were harvested 48 h post-transfection. Supernatants were clarified by sterile filtration (0.22 µm) and cells were scraped from the dished, washed with PBS and collected by centrifugation (500 x *g*, 5min). Cell pellets were lysed in 5 mL lysis buffer (1 % NP-40 / TBS pH 7.5 / 1x cOmplete™ protease inhibitor (Roche, Basel, Switzerland)) for 1 h on ice and cleared by centrifugation (15,000 rpm, 15 min, 4 °C) followed by filtration (0.22 µm).

For immunoprecipitation, lysates and supernatants were incubated with 30 µL NbALFA beads overnight at 4 °C. The beads were subsequently washed extensively with 6 M urea buffer / TBS pH 7.5 followed by TBS pH 7.5 and collected by centrifugation (1,000 × g, 30 s). Bound proteins were resuspended in 30 µL 1× Laemmli buffer, reduced with 5 mM DTT for 30 min at 37 °C, and alkylated with 10 mM iodoacetamide for 30 min at room temperature in the dark. Excess iodoacetamide was quenched by addition of DTT, and ALFA-tagged proteins were eluted by heating at 70 °C for 10 min.

The eluate was divided for downstream analyses: The majority of the sample was separated by SDS-PAGE using a 12% resolving gel and visualized with Protein Detector Stain (BIOZOL, Hamburg, Germany). 2 µL sample was diluted in 1x Lämmli, separated by SDS-PAGE and analyzed by western blot using 0.5 nM NbALFA-IRDye 800CW diluted in 5% non-fat milk in PBS.

### Sample preparation of purified proteins for mass spectrometry

ALFA-positive bands were excised from SDS-PAGE gels and destained with 50% acetonitrile (ACN) in 100 mM ammonium bicarbonate (AmBic) at 37 °C. Gel pieces were dehydrated by incubation with ACN for 2-3 cycles and subsequently washed with 100 mM AmBic. Following rehydration, gel pieces were dehydrated again as described above and incubated with 100 µL of 5 ng/µL sequencing-grade Trypsin Gold (Promega, Madison, WI, USA) in 50 mM AmBic. Proteolytic digestion was performed overnight at 37 °C. Tryptic peptides were sequentially extracted from the gel pieces using 50% acetonitrile / 5% formic acid, 75% acetonitrile / 0.5% formic acid and 100 % ACN. Extracted peptides were dried in a SpeedVac vacuum concentrator (Thermo Savant DNA120 SpeedVac Concentrator; Thermo Fisher Scientific, Waltham, MA, USA) and subjected to a second proteolytic digestion with AspN (Promega, Madison, WI, USA) in 10 mM Tris-HCl (pH 8.0) overnight at 37 °C. Samples were subsequently dried in the SpeedVac vacuum concentrator and reconstituted in 30 µL Orbitrap sample buffer (3 % ACN, 0.1 % TFA).

### Mass spectrometry analysis (LC-MS/MS)

Peptide samples were analyzed on an Orbitrap Fusion Lumos mass spectrometer (Thermo Fisher Scientific, Bremen, Germany) coupled to a nanoLC system (Aurora Ultimate 25×75, ionopticks, Collingwood, Australia) and peptides were separated on a reversed-phase C18 column. MS data were acquired in data-dependent acquisition (DDA) mode. Full MS scans were acquired in the Orbitrap (resolution: 120.000, m/z range: 300-1500), followed by HCD fragmentation (NCE: 35 %) of the most intense precursor ions. MS/MS spectra were recorded via IonTrap with a dynamic exclusion of 15 s. Raw data were processed with FragPipe v24.0 using the plugins MSFragger and IonQuant ((14, 15)) and searched against custom database of IPT repeats encoded by expression plasmids. Trypsin and AspN were specified as proteases, allowing up to one missed cleavage. Carbamidomethylation of cysteine was set as a fixed modification and methionine oxidation as variable. Mono*-*hexose addition (+162.0528 Da) on serine and threonine residues was included as mass offset with neutral loss due to labile modification behavior induced by HCD. Peptide and protein identifications were filtered at a 1% false discovery rate using percolator. Resulting PSMs (see Dataset S3) were validated and extended using Skyline (v26.1, MacCoss Lab, University of Washington). Extracted ion chromatograms (EICs) were generated for peptide signal visualization and evaluation using FreeStyle (Thermo Fisher Scientific).

### Immunofluorescence

2×10^5^ cells were seeded in a six-well on glass coverslips and transfected with the respective plasmid using PEI MAX® 40,000. The cells were washed two times with 1 ml PBS and fixated with 4 % paraformaldehyde in PBS for 10 min at RT. The detection of the ALFA-tag was conducted with a Cy3 conjugated ALFA nanobody (2 nM in 1 % BSA / 0.1 % TritonX-100 / PBS, 1h, RT). Cells were subsequently washed three times with PBS and the cell nuclei were stained with 1 μg/mL Hoechst 33258/PBS for 10 min at RT. After washing the cells twice with PBS and once with water, the glass slides were mounted on with Dako Fluorescence Mounting Medium (Agilent, Santa Clara, USA) on microscope slides. The immunofluorescence samples were analyzed with an Axiovert 200 M microscope (ZEISS, Oberkochen, Germany). The images were processed with the ZEN 2012 software (ZEISS).

### *In silico* alignment of TMEM260 sequons

The amino acid sequences of IPT domains with identified *O-*mannosylation sites were accessed from uniport (Datasheet S1) and aligned with ClustalW from European Molecular Biology Laboratory’s European Bioinformatics Institute (EMBL-EBI, Cambridgeshire, UK).

## Results

### Identification of fibrocystin and fibrocystin-L as *O-*mannosylated proteins in human

The identification of TMEM260 as a protein-specific *O-*mannosyltransferase that selectively modifies extracellular IPT domains (3, 4) prompted us to investigate whether additional IPT-containing proteins might serve as substrates of this pathway. Based on the established substrate specificity of TMEM260 for IPT domains, we therefore considered fibrocystin and fibrocystin-L as candidate substrates, although none of these two proteins had previously been identified as a TMEM260 substrate. To investigate whether fibrocystin and fibrocystin-L are *O-*mannosylated proteins in humans, we first screened publicly available human proteomic datasets for high sequence coverage of each protein using the PeptideAtlas server (13). This search identified a human urinary vesicle proteome dataset (PXD002279) in which fibrocystin was covered by multiple unique peptides and a blood cell surface proteome dataset (PXD005846) providing high sequence coverage of fibrocystin-L. Both datasets were acquired on Q Exactive Orbitrap mass spectrometers using higher-energy collisional dissociation (HCD) for MS2 fragmentation. Since HCD induces fragmentation of *O-*linked glycans, *O-*glycosylated peptide fragments can exhibit a characteristic neutral loss of 162.05 Da corresponding to the loss of a single hexose residue (16). MaxQuant analysis of accessed datasets identified peptide-spectrum matches (PSMs) carrying a +162.05 Da hexose modification in IPT9 of fibrocystin and IPT2 and IPT14 of fibrocystin-L. Thus, the Andromeda algorithm not only identified the mass corresponding to the mannosylated parental ion, but also fragment ions bearing the characteristic neutral loss. This automated analysis was manually confirmed and representative MS2 spectra corresponding to these modified peptides are shown in Fig. 1. As validation of our analysis strategy, we additionally detected *O-*mannosylated peptides from established TMEM260 substrates, including IPT1 of plexin-B2 identified in human primary NK cells and IPT3 of the hepatocyte growth factor receptor (c-MET) identified in the urinary vesicle proteome. All identified PSMs corresponding to hexose-modified peptides derived from IPT domain bearing proteins are listed in Datasheet S2. Together, these results show that fibrocystin and fibrocystin-L are *O-*mannosylated proteins in humans. However, the identification of *O-*mannosylated IPT-derived peptides alone does not establish TMEM260 as the responsible *O-*mannosyltransferase. To determine whether the observed *O-*mannosylation is mediated by TMEM260, we next analyzed fibrocystin and fibrocystin-L in a TMEM260-deficient cellular model.

**Figure 1:**
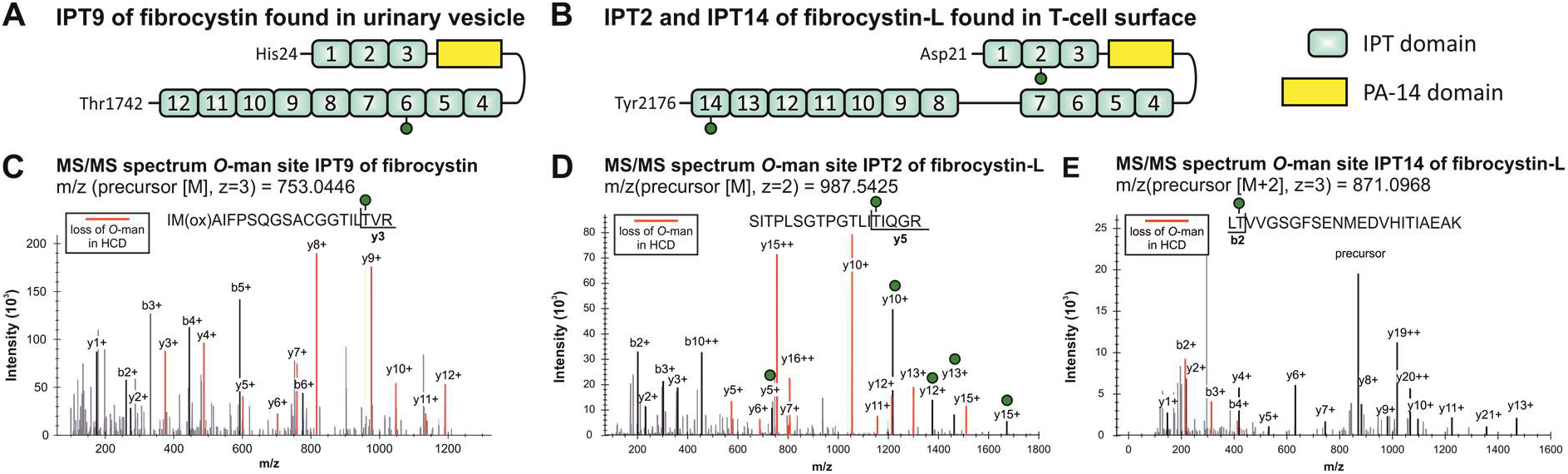
*In silico* identification of *O-*mannosylated IPT domains from fibrocystin and fibrocystin-L in human specimens. **(A,B)** Extracellular domain organization of IPT domains reported for fibrocystin **(A)** or fibrocystin-L **(B)** (7). IPT domains are shown in light green, the PA-14 domain in yellow, and *O-*mannosylated IPT domains are labeled with green circles. Publicly available proteomic datasets from human urinary extracellular vesicles and primary blood cells were retrieved from PRIDE and were screened for *O-*mannosylated IPT-derived peptides. **(C-E)** MS/MS spectra of *O-*mannosylated peptides derived from fibrocystin IPT9 **(C)**, fibrocystin-L IPT2 **(D)**, and fibrocystin-L IPT14 with annotated fragments **(E)**. Characteristic neutral losses of -162.05 Da are shown in red and +162.05 Da hexose modified fragments are labeled with a green circle.

### TMEM260 complementation restores *O-*mannosylation of fibrocystin and fibrocystin-L IPT repeats in TMEM260-KO cells

Using CRISPR-Cas9 genome editing, we targeted exon 3 of the TMEM260 locus in HEK293 cells and established a TMEM260-KO cell line. Since HEK293 cells lack endogenous *PKHD1* and *PKHD1L1* expression, we generated secreted recombinant IPT repeats comprising three to four IPT domains. IPT repeats were *C*-terminally fused to an ALFA-His10 tag to enable detection, enrichment, and subsequent MS analysis (Fig. 2A). Although this approach does not fully recapitulate the native protein context, it enables systematic analysis of TMEM260 substrate recognition, which is mediated by primary protein sequence features.

**Figure 2:**
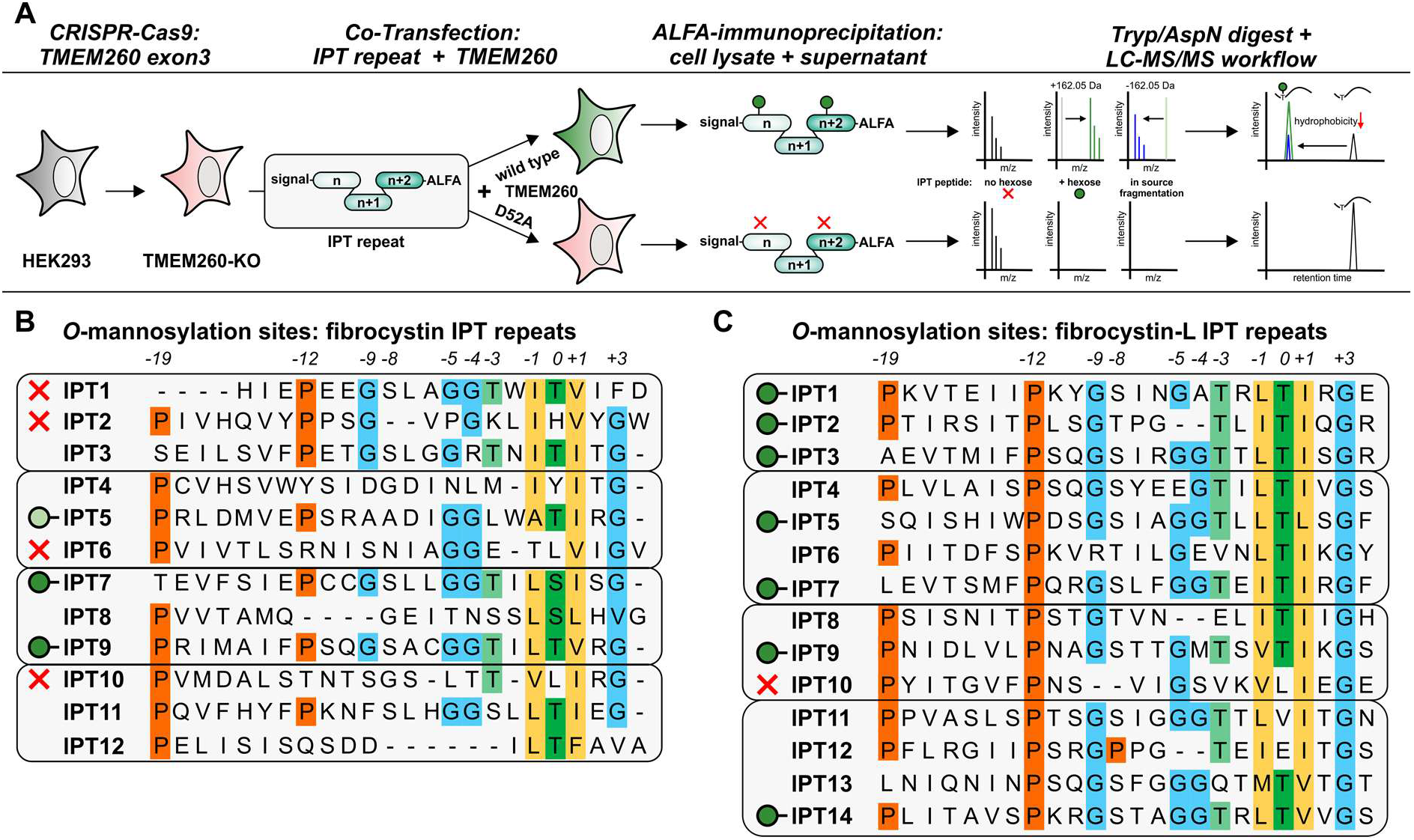
TMEM260 complementation restores site-specific *O-*mannosylation of conserved fibrocystin and fibrocystin-L sequons in TMEM260-KO cells. **(A)** Experimental workflow for the identification of TMEM260-mediated *O-*mannosylation. TMEM260-KO cells were co*-*transfected with secreted IPT repeat constructs and either wild-type TMEM260 or the catalytically inactive TMEM260(D52A) mutant. IPT repeats were enriched by ALFA-tag immunoprecipitation from cell lysates or conditioned medium, and *O-*mannosylation was analyzed by LC-MS/MS. *O-*mannosylated peptides (green) displayed characteristic retention time shifts and in-source fragmentation behavior, resulting in co-elution of unmodified peptide species (blue). Sequence alignment of conserved TMEM260 sequon sequences from fibrocystin **(B)** and fibrocystin-L **(C)**. *O-*mannosylated sequons are indicated by green circles, weakly *O-*mannosylated sequons by light green circles, and sequons for which no *O-*mannosylation was detected by red crosses. Sequons that were not identified in the LC-MS/MS dataset are left unlabeled.

To assess TMEM260-mediated *O-*mannosylation of fibrocystin and fibrocystin-L, IPT repeats were co*-*expressed in TMEM260-KO cells with either wild-type TMEM260 (WT) or a catalytically inactive TMEM260 mutant (D52A) and purified from cell lysate and conditioned media (Fig. 2A). Both TMEM260 variants exhibited comparable expression levels and ER-like localization, indicating that the D52A mutation did not impair protein expression or targeting (Fig. S1). An IPT repeat construct comprising IPT1-3 of plexin-D1 was included as a positive control. Despite variable secretion levels, sufficient material was obtained by enrichment from whole-cell lysates among individual IPT repeats (Fig. S2-4), hence, the LC-MS/MS analysis was focused on lysate-derived samples. Notably, secretion of IPT11-14 from fibrocystin-L and IPT1-3 from plexin-D1 was only observed upon complementation with WT TMEM260, suggesting that TMEM260-mediated *O-*mannosylation may influence secretion efficiency of selected IPT repeats.

We validated our LC-MS/MS analysis strategy by confirming *O-*mannosylation of the plexin-D1 IPT1-3 repeat. Hexose-modified peptides were detected for all three IPT domains upon complementation with WT TMEM260, but not with the inactive TMEM260 D52A mutant (Fig. S5). Applying this validated approach to fibrocystin and fibrocystin-L IPT repeats, we identified additional TMEM260-mediated hexose modifications beyond the IPT domains detected in human proteomic datasets (Fig. 2B, Fig. S5). These included IPT7 of fibrocystin and IPT1, IPT3, IPT5, IPT7, and IPT9 of fibrocystin-L. For IPT5 of fibrocystin, only a minor fraction of the detected peptide (∼10%) carried the hexose modification, suggesting limited substrate recognition at this site. Interestingly, for some peptide species, the chromatographic signal corresponding to the *O-*mannosylated peptide was associated with MS spectra containing m/z signals for both the *O-*mannosylated and the corresponding unmodified peptide. Thus, although the peptide eluted at the retention time expected for the *O-*mannosylated species, the MS analysis revealed a co-occurring signal corresponding to the unmodified peptide. This observation is consistent with partial in-source fragmentation of the glycosidic bond during ionization, resulting in neutral loss of the *O-*mannosyl group from a fraction of the precursor ions prior to MS1 and MS2 acquisition (Fig. 2A, Fig. S5).

### TMEM260-mediated *O-*mannosylation accommodates threonine-to-serine substitution in an IPT-derived reporter lacking the *N*-terminal sequon context

The newly identified *O-*mannosylation sites within the IPT domains of fibrocystin and fibrocystin-L provide additional evidence for flexibility in the sequence requirements of TMEM260 substrate recognition (Fig. 2, Fig. S6). These sites deviate from the previously defined TMEM260 sequon (4), suggesting that the sequence constraints governing substrate recognition may be broader than previously appreciated. In detail, variations at conserved position −8 (proline) as well as *N*-terminal truncations of the motif in fibrocystin-L IPT2 were tolerated by TMEM260.

Most strikingly, TMEM260 also recognizes and glycosylates serine at position 0 within IPT7 of fibrocystin, suggesting that the canonical threonine residue is not strictly required for TMEM260-mediated *O-*mannosylation. Prompted by this unexpected observation, we asked whether this apparent flexibility in acceptor residue identity also applies to a distinct TMEM260 substrate. We therefore turned to plexin-B2, a well-established TMEM260 substrate, and analyzed *O-*mannosylation of reporter constructs in which the plexin-B2 IPT1 *O-*mannosylation motif was fused to the *N-*terminus of secreted enhanced green fluorescent protein (eGFP). The reporter panel consisted of a construct containing the intact plexin-B2 IPT1 sequon (positions −19 to +3), a variant with an *N-*terminal truncation of the motif (positions −12 to +3), and a corresponding truncated variant in which the threonine residue at position 0 was replaced by serine. All reporter constructs were enriched from cell lysates (Fig. S4) and analyzed by mass spectrometry (Fig. 3, Fig. S5). Strikingly, neither *N-*terminal shortening of the corresponding sequence context nor *N-*terminal shortening together with replacement of the canonical threonine by serine abrogated *O-*mannosylation at position 0. The *N-*terminally truncated variants displayed reduced modification efficiency compared with the intact reporter, indicating that residues outside the core sequon contribute to efficient TMEM260 substrate recognition but are not essential for catalysis. Taken together, these findings demonstrate that TMEM260 can accommodate serine as an *O-*mannosylation acceptor not only in IPT7 of fibrocystin, but also in an unrelated substrate context of plexin-B2.

**Figure 3:**
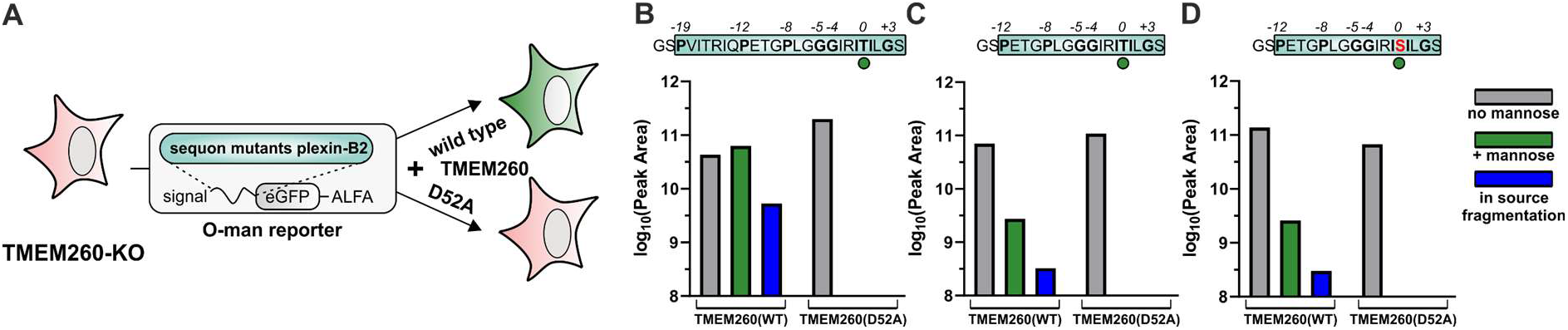
Threonine and serine serve as acceptor residues for TMEM260-mediated *O*-mannosylation in *N*-terminally truncated reporters. **(A)** Schematic representation of the secreted eGFP reporter constructs carrying the plexin-B2 IPT1 *O*-mannosylation sequon fused to the *N*-terminus of eGFP. Reporter variants contained either the intact sequon, an *N*-terminal truncation, or an *N*-terminally truncated sequon carrying a threonine-to-serine substitution at position 0. Reporters were co-expressed with TMEM260 and enriched via the *C*-terminal ALFA tag prior to mass spectrometry analysis. **(B–D)** Integrated peak area of signals corresponding to m/z values of unmodified (grey), *O*-mannosylated (green), and in-source fragmented *O*-mannosylated (blue) peptides determined by LC-MS/MS. Reporter peptide sequences are shown above each graph, with highlighted conserved residues.

### The ARPKD-associated fibrocystin variant L1407R abolishes *O-*mannosylation of IPT9

Despite the apparent heterogeneity of the TMEM260 sequon, predominantly hydrophobic, nonpolar amino acids (Ile, Leu, Val Gly) at positions −1 and +1 as well as glycine residues at position -4 and +3 remained conserved in all *O-*mannosylated IPT domains, suggesting that these residues represent core determinants for substrate recognition. To establish a potential link between TMEM260-mediated *O-*mannosylation of fibrocystin and its associated diseases, we investigated the ARPKD-associated fibrocystin variant L1407R (17). The L1407R substitution which is located in IPT9 replaces a hydrophobic leucine with a positively charged, highly polar arginine residue at position -1 of the TMEM260 recognition motif.

We encountered an analytical challenge with the L1407R variant, as the mutation introduces an additional tryptic cleavage site, resulting in a peptide that was too short to be reliably detected by our conventional LC–MS/MS workflow. However, the corresponding unmodified miscleaved peptide (TVRGLLLNSR) was readily detected by MS1 and its identity was confirmed by MS2 fragmentation. In contrast, the corresponding *O-*mannosylated species was not detected (Fig S5G). These findings are consistent with the loss of TMEM260-mediated *O-*mannosylation caused by the L1407R substitution and demonstrate that the conserved unpolar residue at the -1 position is critical for TMEM260 substrate recognition.

## Discussion

TMEM260 has recently been identified as a protein-specific *O-*mannosyltransferase that catalyzes the transfer of a single mannose residue to threonine residues within extracellular immunoglobulin, plexin, transcription factor (IPT) domains. Larsen et al. demonstrated that TMEM260 selectively *O-*mannosylates IPT domains of a defined subset of receptor proteins, including the semaphorin receptors plexins as well as the receptor tyrosine kinases cMET and RON, thereby establishing a distinct mammalian *O-*mannosylation pathway. TMEM260-mediated *O-*mannosylation was shown to be important for the maturation and trafficking of several of these receptors, linking this modification to receptor function and developmental processes.(3)

More recently, structural analyses of human TMEM260 in complex with an acceptor peptide derived from plexin-B2 revealed the molecular basis of substrate recognition and identified a conserved *O-*mannosylation sequon that contributes to acceptor specificity. These findings established a defined sequence context for TMEM260-mediated *O-*mannosylation and suggested that substrate recognition involves both the modified residue and its surrounding sequence environment (4).

Our identification of fibrocystin and fibrocystin-L as additional TMEM260-dependent *O-*mannosylated proteins expands the known substrate spectrum of this enzyme and, importantly, reveals that the sequence requirements for TMEM260-mediated *O-*mannosylation may be more flexible than previously appreciated. In addition, our findings demonstrate that TMEM260-mediated *O-*mannosylation promotes the secretion of protein fragments containing IPT domains, supporting a functional role for this modification in protein trafficking. The identification of fibrocystin as a TMEM260 substrate further establishes a direct molecular link between TMEM260-mediated *O-*mannosylation and ARPKD, providing a potential mechanistic connection between TMEM260 dysfunction and renal disease.

Beyond expanding the biological spectrum of TMEM260 substrates, our findings redefine the sequence requirements governing TMEM260-mediated *O-*mannosylation. The newly identified *O-*mannosylation sites reveal that TMEM260 tolerates considerable variation within the *N-*terminal region of the previously described sequon (4). Particularly, IPT2 of fibrocystin-L lacks residues within the *N-*terminal portion of the sequon while remaining efficiently *O-*mannosylated. These observations suggest that the luminal region of TMEM260 provides a flexible substrate-binding platform that accommodates sequence variability, whereas conserved residues surrounding the acceptor site anchor the substrate within the catalytic center. Consistent with this model, TMEM260-mediated *O-*mannosylation was retained in a reporter construct containing an *N*-terminally truncated sequon, further supporting that the full context of a conserved *N*-terminal sequon is not required for TMEM260 activity. Our data further expand the substrate specificity of TMEM260 by demonstrating that serine can serve as an alternative acceptor residue for *O-*mannosylation. Although serine modification had been proposed as a possible feature of TMEM260 substrates (3, 4), it has not been demonstrated before. In contrast, our findings indicate that specific residues surrounding the glycosylation site impose stronger constraints on substrate recognition. The conserved glycine at position +3 appears particularly important, potentially reflecting structural requirements of small residues like glycine within the catalytic pocket (4). Notably, for IPT1 of fibrocystin no *O-*mannosylation was measured potentially caused by the presence of bulky phenylalanine at position +3.

The importance of conserved residues surrounding the *O-*mannosylation site was further supported by the analysis of fibrocystin L1407R which is a sequence variant associated with ARPKD. In IPT9 of fibrocystin, substitution of the conserved -1 leucine (L1407) with arginine resulted in loss of detectable *O-*mannosylation. All *O*-mannosylated IPT domains of fibrocystin or fibrocystin-L identified in this study contain either leucine, isoleucine or valine at position -1 with the exception of fibrocystin IPT5 containing alanine. These findings indicate that a hydrophobic residue at this position contributes to TMEM260 substrate recognition. However, the higher modification efficiency observed with leucine, isoleucine and valine compared with alanine in IPT5 suggests that substrate recognition might be influenced by the hydrophobicity of the respective amino acid at this position which is considerable higher for isoleucine, valine and leucine compared to alanine. Accordingly, substitution with the positively charged arginine as in the L1407R variant completely abolished *O-*mannosylation, further supporting a preference for hydrophobic residues at this position while highlighting the contribution of residue-specific features to TMEM260 substrate recognition. This may reflect differences in hydrophobic interactions between the substrate peptide and the TMEM260 catalytic center, where larger hydrophobic side chains, such as leucine, isoleucine, or valine, may establish more favorable hydrophobic interactions than alanine.

The molecular mechanisms connecting loss of TMEM260 function to SHDRA phenotypes remain incompletely understood. The link between TMEM260 deficiency and the cardiac manifestations of SHDRA, particularly persistent truncus arteriosus, has previously been suggested by the identification of plexin-D1 as a TMEM260 substrate, providing a mechanistic connection between impaired *O-*mannosylation and the cardiac phenotype (3, 6, 18). In contrast, the molecular basis of the renal manifestations of SHDRA has remained elusive. The identification of fibrocystin as a TMEM260 substrate now provides a direct molecular link between TMEM260 deficiency and renal disease, as fibrocystin is a causative gene underlying autosomal recessive polycystic kidney disease (ARPKD). Consistent with this disease association, the ARPKD-associated L1407R missense variant within the TMEM260-modified IPT9 domain abolished detectable *O-*mannosylation, highlighting the functional relevance of fibrocystin *O-*mannosylation.

The patient-specific fibrocystin missense variant Leu1407Arg (L1407R) is affecting a potential *O*-mannosylation site in the IPT9 of fibrocystin and has been identified in patients with ARPKD (8, 9, 19). This mutation is currently classified in ClinVar database (accession: VCV000813383.13) as pathogenic/likely pathogenic (20). Notably, L1407R was identified in trans with a truncating variant in a patient who survived the neonatal period but subsequently developed renal manifestations of ARPKD (8). In contrast, patients carrying two truncating *PKHD1* variants generally present with a severe phenotype, often resulting in perinatal or neonatal death, whereas heterozygous carriers are typically clinically unaffected (21). The comparatively milder clinical course of the patient carrying one truncating allele and the L1407R allele is therefore consistent with the possibility that L1407R retains residual fibrocystin function. Nevertheless, the L1407R allele was insufficient to prevent disease in the presence of a truncating allele, as would be expected for a fully functional second allele. Given that L1407R alters a residue within a potential *O*-mannosylation site, the substitution may additionally affect fibrocystin *O*-mannosylation and thereby contribute to the molecular consequences of the variant.

Interestingly, *PKHD1L1* variants have been associated with hearing loss, suggesting a potential role of fibrocystin-L in the development the auditory system (22, 23). Notably, auditory abnormalities have also been reported in a subset of individuals with SHDRA syndrome (5, 24), raising the possibility that impaired *O-*mannosylation of fibrocystin-L contributes to auditory defects associated with TMEM260 deficiency.

The biological function of IPT domain *O-*mannosylation and its functional relevance for the modified proteins remain largely unresolved. Previous studies have demonstrated that ER-derived *O-*mannosylation can influence protein folding, receptor maturation, and trafficking (3). Consistent with this concept, we observed that TMEM260-mediated *O-*mannosylation enable secretion of overexpressed IPT repeats from fibrocystin-L (IPT11–14) and plexin-D1 (IPT1–3). These findings suggest that *O-*mannosylation may contribute to the maturation or structural stabilization of IPT-containing extracellular regions. Given the exceptionally large extracellular regions of fibrocystin and fibrocystin-L, comprising numerous IPT domains, *O-*mannosylation may represent an important mechanism to maintain the extracellular structural integrity.

Taken together, our findings demonstrate that TMEM260 recognizes a broader and more diverse spectrum of IPT domain substrates than previously appreciated. We identify fibrocystin and fibrocystin-L as *O-*mannosylated proteins, providing a potential molecular link between TMEM260 dysfunction and *PKHD1*-associated disease phenotypes in SHDRA syndrome. As the TMEM260 recognition sequon is more degenerated than previously assumed, additional proteins may represent yet unidentified TMEM260 substrates.

## Acknowledgements

This study was supported by the Deutsche Forschungsgemeinschaft (DFG, German Research Foundation) for FOR2953 (project: 409784463 for FFRB), FOR2509 (project: 289991887, for FFRB). We thank Birgit Tiemann for her contribution to the generation of the nanobody tools and for providing protocols for the production and purification of recombinant proteins. We also thank Laura Sophie Dräger for her valuable support in the evaluation of MS data.

## Declaration of generative AI and AI-assisted technologies in the writing process

The authors used ChatGPT (OpenAI) during the preparation of this manuscript to improve language and readability. The authors reviewed and edited the output as necessary and take full responsibility for the content of the publication.

## Author contributions

Conceptualization: FFRB, JK, SBM. Methodology: JK, CH. Investigation: JK, CH, CG, AP. Supervision, resources, and project administration: FFRB. Visualization: JK. Writing – original draft: JK, FFRB. Writing – review and editing: all authors contributed to and approved the final manuscript.

## Supplemental data

This article contains supplemental data.

