## Supplementary information for "*O*-mannosylation of fibrocystin and fibrocystin-L expands TMEM260 sequon selectivity and provides a potential link to the renal and auditory phenotypes of SHDRA"

*Corresponding author:

Falk F. R. Buettner

**Content of supplement:**

Table S1

Figure S1, Figure S2, Figure S3, Figure S4, Figure S5, Figure S6

**Additional supplementary material:**

Supplementary Datasets containing:

Dataset S1: List of human proteins bearing IPT domains annotated in uniprot database

Dataset S2: List of hexose modified IPT domain derived peptides found in human specimens

Dataset S3: List of hexose (non-) modified peptides derived from recombinant IPT repeats found in fragpipe search

**Table S1: Plasmids used in this study.** Protein residues encoded by expression plasmids are displayed and corresponding accessions are shown in italic (*uniprot*) or with preceding # (addgene). The molecular weight ($M_W$) corresponds to the mature protein.

| plasmid / accession | backbone | Encoded protein / RNA | $M_W$ / kDa | mutations |
|---|---|---|---|---|
| hPlexin-D1$^{IPT1-3}$ / *Q9Y4D7* | pcDNA3.1 | Signal(C6)-Leu$^{868}$-Lys$^{1182}$-ALFA-His10 | 37.3 | *H. sapiens* codon optimized |
| hTMEM260 / *Q9NX78* | pcDNA3.1 | Met$^{1}$- Val$^{707}$-ALFA | 81.3 | Native coding sequence |
| hTMEM260(D52A) | | | 81.3 | Asp$^{52}$ (GAC) → Ala (GCC) |
| hPKHD1$^{IPT1-3}$ / *P08F94* | pcDNA3.1 | Signal(C6)-His$^{24}$-Gly$^{335}$-ALFA-His10 | 37.8 | Native coding sequence |
| hPKHD1$^{IPT4-6}$ / *P08F94* | pcDNA3.1 | Signal(C6)-Pro$^{931}$-Ile$^{1192}$-ALFA-His10 | 31.6 | Native coding sequence |
| hPKHD1$^{IPT7-9}$ / *P08F94* | pcDNA3.1 | Signal(C6)-Thr$^{1196}$-Arg$^{1481}$-ALFA-His10 | 33.6 | Native coding sequence |
| hPKHD1$^{IPT7-9}$(L1407R) | | | 33.6 | Leu$^{1407}$ (CTT) → Arg (CGT) |
| hPKHD1$^{IPT10-12}$ / *P08F94* | pcDNA3.1 | Signal(C6)-Pro$^{1486}$-Arg$^{1742}$-ALFA-His10 | 31.3 | Native coding sequence |
| hPKHD1L1$^{IPT1-3}$/ *Q86WI1* | pcDNA3.1 | Signal(C6)-Ser$^{23}$-Arg$^{364}$-ALFA-His10 | 41.2 | Native coding sequence |
| hPKHD1L1$^{IPT4-7}$/ *Q86WI1* | pcDNA3.1 | Signal(C6)-Pro$^{1067}$-Ser$^{1410}$-ALFA-His10 | 40.4 | Native coding sequence |
| hPKHD1L1$^{IPT8-10}$/ *Q86WI1* | pcDNA3.1 | Signal(C6)-Pro$^{1566}$-Val$^{1828}$-ALFA-His10 | 31.2 | Native coding sequence |
| hPKHD1L1$^{IPT11-14}$/ *Q86WI1* | pcDNA3.1 | Signal(C6)-Pro$^{1831}$-Tyr$^{2176}$-ALFA-His10 | 41.2 | Native coding sequence |
| *O*-man reporter long (T): hPlexin-B2$^{IPT1}$/ *O15031* | pcDNA3.1 | Signal(C6)-Pro$^{803}$-Ser$^{826}$-eGFP-ALFA-His10 | 32.7 | *H. sapiens* codon optimized |
| *O*-man reporter short (T): hPlexin-B2$^{IPT1}$/ *O15031* | pcDNA3.1 | Signal(C6)-Pro$^{810}$-Ser$^{826}$-eGFP-ALFA-His10 | 31.9 | *H. sapiens* codon optimized |
| *O*-man reporter short (S): hPlexin-B2$^{IPT1}$/ *O15031* | pcDNA3.1 | Signal(C6)-Pro$^{810}$-Ser$^{826}$-eGFP-ALFA-His10 | 31.9 | *H. sapiens* codon optimized Thr$^{822}$ (ACC) → Ser (TCC) |
| pCas9-GFP / #44719 | pCAG | Cas9-P2A-eGFP (1) | N/A | N/A |
| sgRNA plasmid / #68463 | N/A | gRNA U6 promotor-controlled expression (2) | N/A | N/A |

Signal(C6): Signal peptide from human complement component C6 with GS linker (MARRSVLYFILLNALINKGQA-GS, *P13671*), ALFA: extended ALFA-tag (PSRLEEELRRRLTEPTG) (3) , His10: ten histidine residues (HHHHHHHHHH), eGFP: enhanced green fluorescent protein

**Table S2: Genotype of validated HEK293 TMEM260 KO clone and guide RNA sequence (gRNA) used for CRISPR-Cas9 mediated KO generation.**

| clone | gRNA sequence (5' to 3' - *PAM*) | locus | Indel | protein sequence at gRNA site |
|---|---|---|---|---|
| TMEM260-KO | ACAGAGAAGATTGACGCGGT- *AGG* | Exon3 (-) strand | -7bp | G$^{88}$-S-I-A-S-…-L-E-Q$^{101}$-**stop** |
| HEK293 WT | | | WT | G$^{88}$-S-I-A-Y-R-V-N-L$^{96}$ |

Reference mRNA sequence used: NM_017799.4 (NCBI database)

**Table S3: List of DNA oligos used for PCR / cloning procedures.**

| name | primer sequence (5' to 3') | template | purpose |
|---|---|---|---|
| $P_{fw}$(D52A) | ACCGGGGGGAGCCTCCGGGGAACTGATCAC | *hTMEM260* mRNA (NM_017799.4) | Site directed mutagenesis |
| $P_{rev}$(D52A) | GTTCCCCGGAGGCTCCCCCCGGTACCGAAG | | |
| $P_{fw}$(I911L) | GCTTACAATGAGAGGCCGGAATCTCGGACGCCGATTATCC | Codon optimized hPlexin-D1 coding sequence | Site directed mutagenesis |
| $P_{rev}$(I911L) | CCGGCCTCTCATTGTAAGCAACGTGCCGCCATCCAGTGG | | |
| $P_{fw}$(L1407R) | CGCAGGGTTCGGCATGTGGTGGGACCATACGTACT | *hPKHD1* mRNA (NM_138694.4) | Site directed mutagenesis |
| $P_{rev}$(L1407R) | CTGGAGTTAAGAAGCAACCCCCTCACAGTACGTATG | | |
| $P_{fw}$(gDNA) | GATAGCAGTGTGGTCTTTCA | *hTMEM260* locus (NC_000014.9: 56579525-56663165) | gDNA screening of TMEM260 KO clones |
| $P_{rev}$(gDNA) | CTACATCTCCAACTGCTCAA | | |
| $P_{fw}$(cDNA) | AGACTCCGGGGAACTGAT | *hTMEM260* mRNA (NM_017799.4) | cDNA screening of TMEM260 KO clones |
| $P_{rev}$(cDNA) | TCCAGAATCGTTCCACCACAC | | |

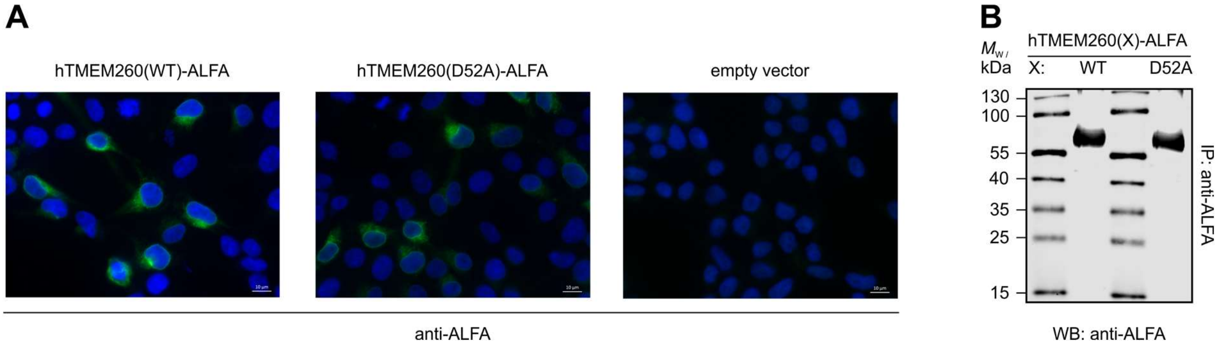


**Figure S1: Wild-type hTMEM260 and the inactive D52A mutant show similar expression levels and subcellular localization. (A)** Immunofluorescence microscopy of HEK293 cells overexpressing ALFA-tagged TMEM260(WT) or TMEM260(D52A), detected using a Cy3-coupled anti-ALFA nanobody (green). Nuclei were stained with Hoechst 33342 (blue). **(B)** Anti-ALFA Western blot of ALFA immunoprecipitants from HEK293 cells overexpressing TMEM260(WT)-ALFA or hTMEM260(D52A)-ALFA.

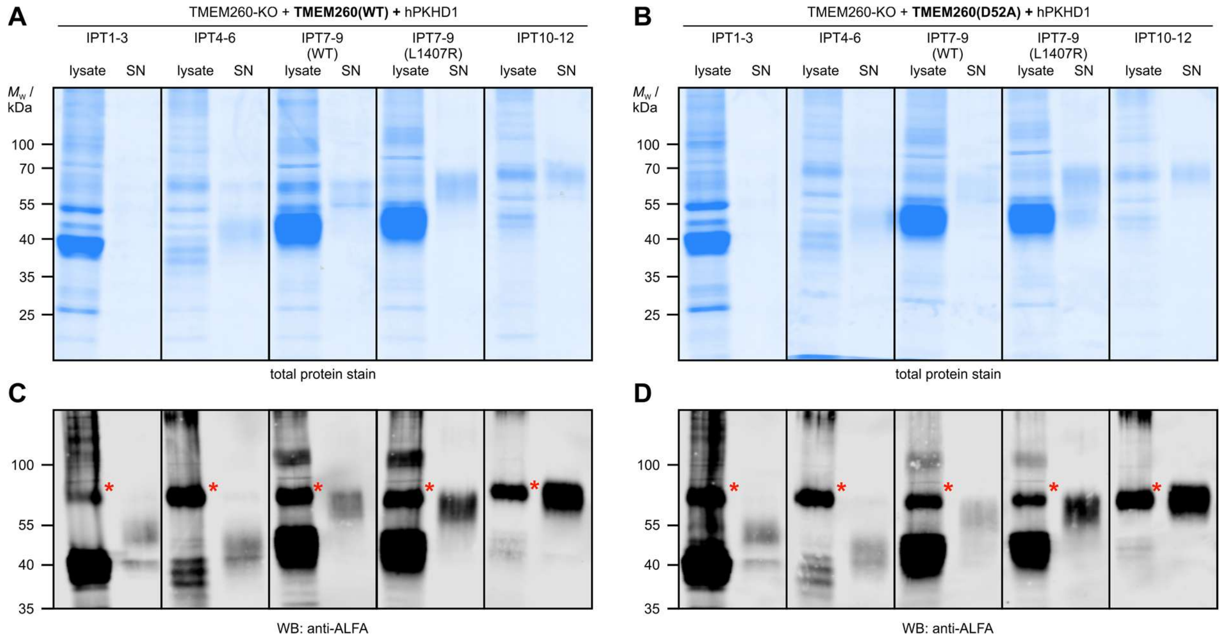


**Figure S2: Expression and purification of recombinant fibrocystin IPT tandem constructs in TMEM260-KO cells complemented with wild-type or TMEM260(D52A).** HEK293 TMEM260 knockout cells were complemented with either TMEM260(WT)-ALFA **(A,C)** or the catalytically impaired TMEM260(D52A)-ALFA variant **(B,D)** and co-expressed with hPKHD1-derived IPT repeat constructs. Recombinant IPT repeats comprising IPT domains 1-3, 4-6, 7-9, or 10-12 were expressed as soluble ALFA-His10-tagged proteins. **(A,B)** Total protein stained SDS-PAGEs of anti-ALFA immunoprecipitations from lysates and cell culture supernatant (SN) of transfected cells. **(C,D)** Western blot analysis of corresponding samples using an anti-ALFA nanobody to detect protein constructs. Red asterisks label ALFA-positive signal at the expected molecular weight of co-expressed TMEM260-ALFA variants in Western blot. IPT7–9(L1407R) represents the indicated IPT repeat from PKHD1 ARPKD-variant.

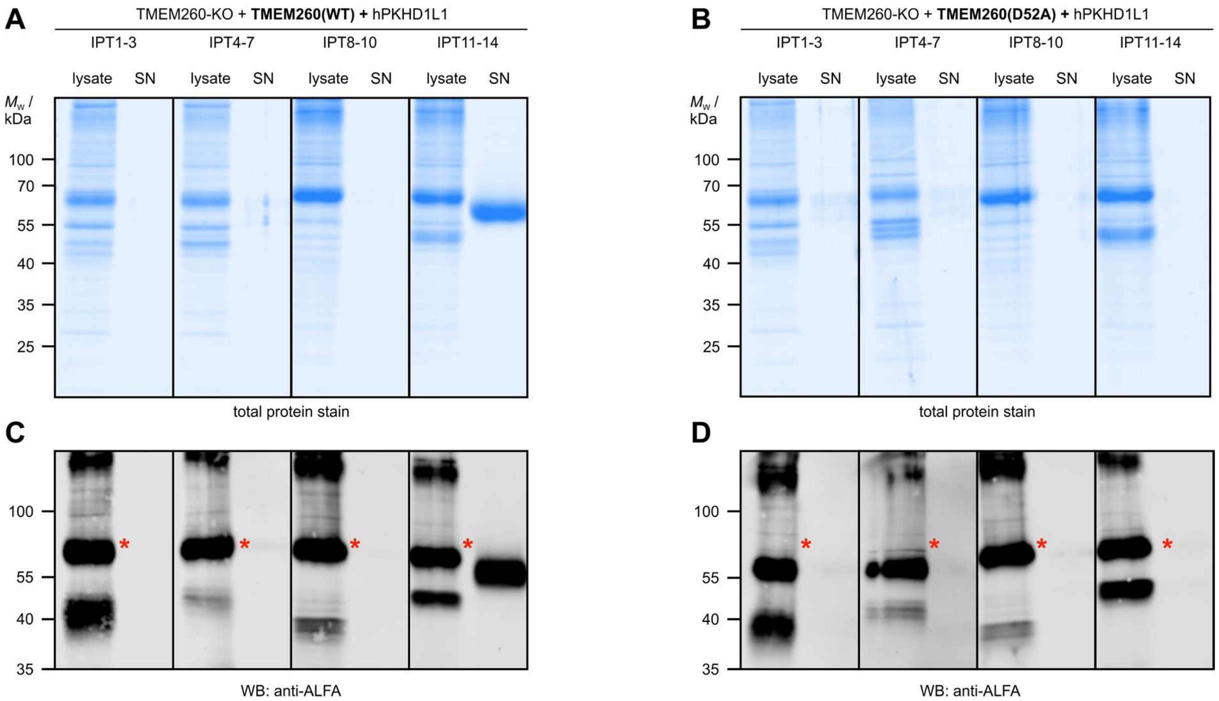


**Figure S3: Expression and purification of recombinant fibrocystin-L IPT tandem constructs in TMEM260-KO cells complemented with wild-type or TMEM260(D52A).** HEK293 TMEM260 knockout cells were complemented with either TMEM260(WT)-ALFA **(A,C)** or the catalytically impaired TMEM260(D52A)-ALFA variant **(B,D)** and co-expressed with hPKHD1-derived IPT repeat constructs. Recombinant IPT repeats comprising IPT domains 1-3, 4-7, 8-10, or 11-14 were expressed as soluble ALFA-His10-tagged proteins. **(A,B)** Total protein stained SDS-PAGEs of anti-ALFA immunoprecipitations from lysates and cell culture supernatant (SN) of transfected cells. **(C,D)** Western blot analysis of corresponding samples using an anti-ALFA nanobody to detect protein constructs. Red asterisks label ALFA-positive signal at the expected molecular weight of co-expressed TMEM260-ALFA variants in Western blot.

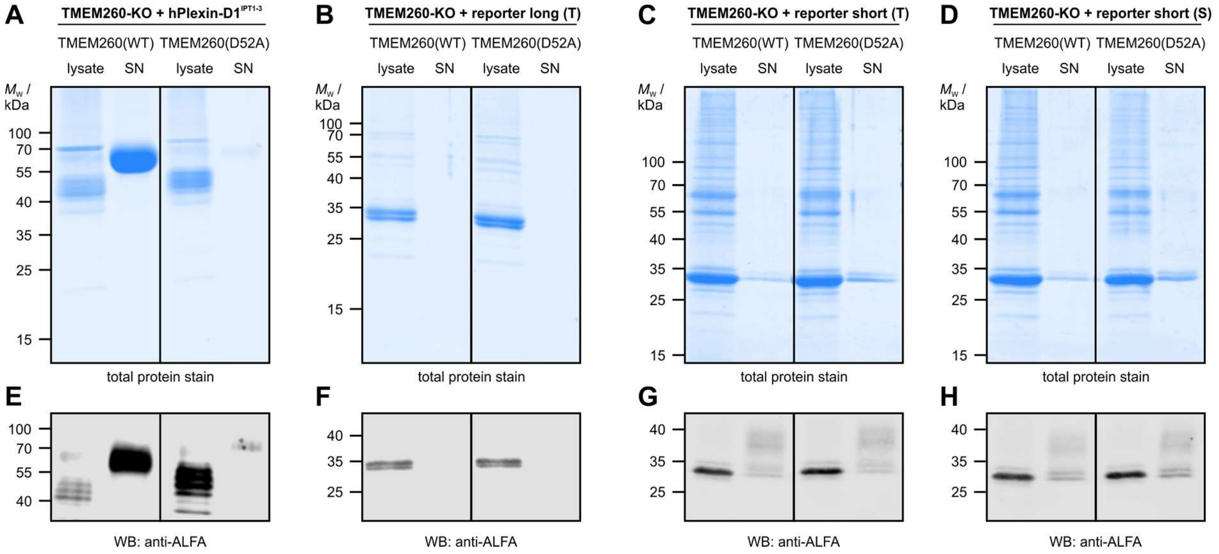


**Figure S4: Expression and purification of recombinant plexin-D1 IPT tandem and *O*-mannosylation reporter constructs in TMEM260-KO cells complemented with wild-type or TMEM260(D52A).** HEK293 TMEM260 knockout cells were complemented with either TMEM260(WT)-ALFA or the catalytically impaired TMEM260(D52A)-ALFA variant and co-expressed with constructs. Recombinant IPT repeats comprising IPT domains 1-3 from plexin-D1 and reporter constructs were expressed as soluble ALFA-His10-tagged proteins. **(A-D)** Total protein stained SDS-PAGEs of anti-ALFA immunoprecipitations from lysates and cell culture supernatant (SN) of transfected cells. **(E-H)** Western blot analysis of corresponding samples using an anti-ALFA nanobody to detect protein constructs.

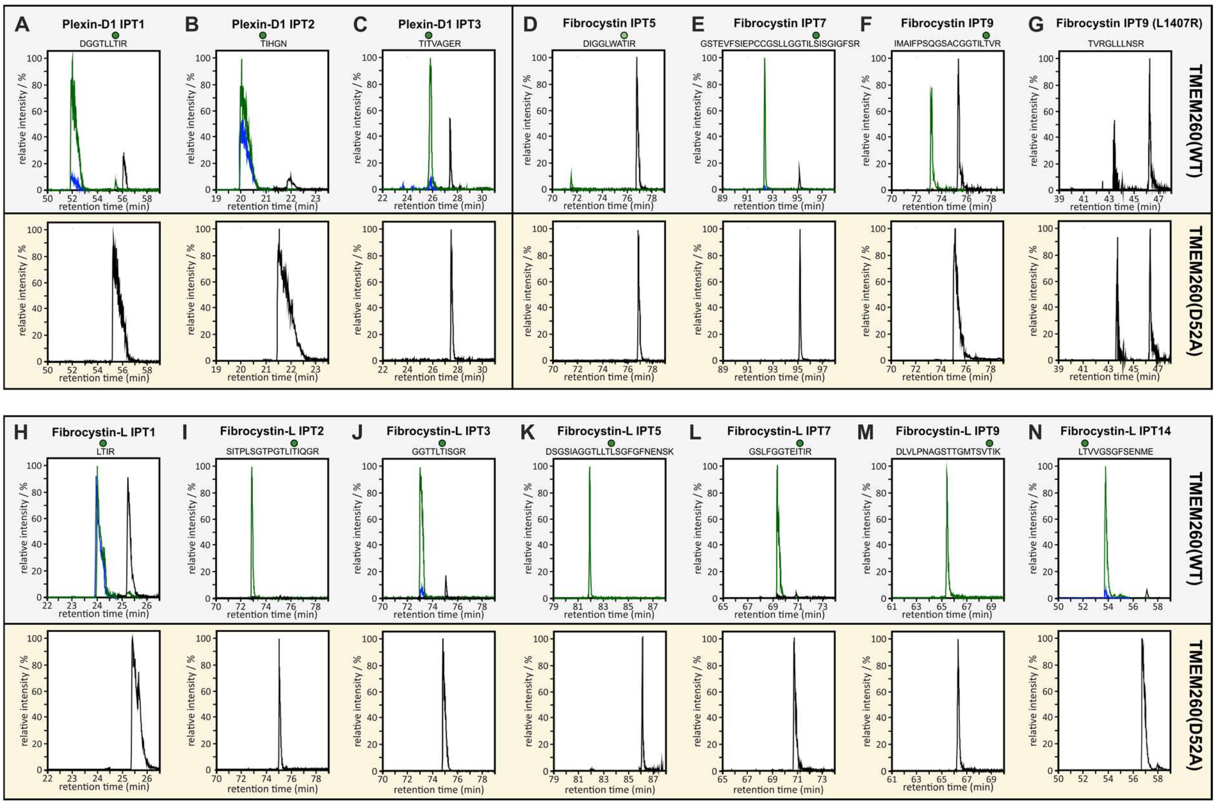


**Figure S5: Extracted ion chromatograms (EICs) demonstrating TMEM260-dependent *O*-mannosylation of IPT repeats.** EICs corresponding to m/z values of IPT domain-derived peptides from TMEM260-KO cells complemented with either TMEM260(WT) (gray) or the catalytically inactive TMEM260(D52A) mutant (yellow) are shown within selected retention time windows. Signals corresponding to non-modified peptides are indicated in black, whereas hexose-modified peptides (+162.05 Da) are shown in green. Due to in-source fragmentation of *O*-mannosylated peptides, non-modified peptide ions can be detected at the same retention times as hexose-modified peptides (blue). Signal intensities were normalized to the highest intensity observed within each individual EIC. The corresponding IPT domains and *O*-mannosylated peptides are indicated above each EIC trace. All EICs were generated from peptide masses whose identities were validated by MS/MS spectra.

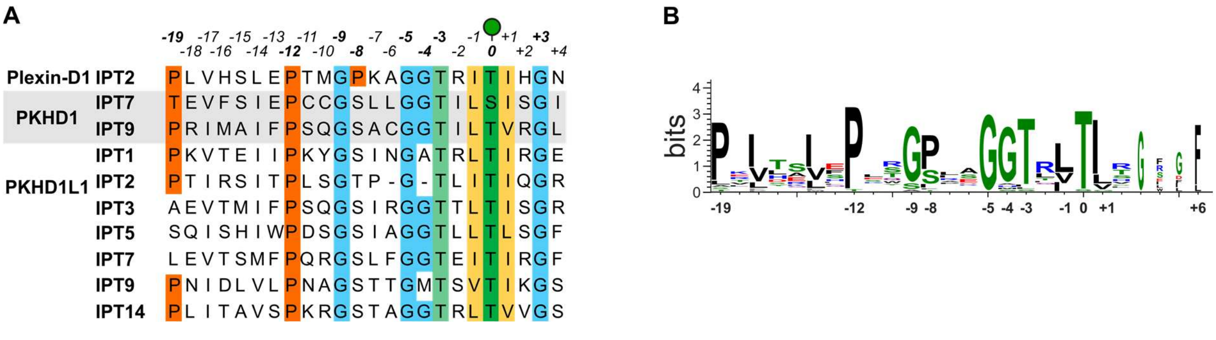


**Figure S6: Sequence alignment of TMEM260-mediated *O*-mannosylated IPT domains reveals conserved sequence features surrounding the modification site. (A)** Alignment of all experimentally identified *O*-mannosylation sites found in human fibrocystin (*PKHD1*) and fibrocystin-L (*PKHD1L1*) displayed similarly as previously shown for plexins, MET and RON by Cifuente *et al*. (4). Amino acids are numbered relative to the modified residue at position 0, and conserved residues are highlighted. The *O*-mannosylated amino acid is indicated in dark green, while conserved residues within the TMEM260 recognition motif are indicated by colored shading. **(B)** Sequence logo generated from all known TMEM260-mediated *O*-mannosylated IPT sequences present in plexins, MET and RON (cifuente *et al.* (4)) as well as fibrocystin and fibrocystin-L (this study). This combined analysis shows conserved amino acid preferences surrounding the modification site. IPT5 of PKHD1 is not considered in this alignment to due less pronounced degree of *O*-mannosylation at this site.
